# Cryptic connectivity between hyporheic and riparian zones via winged aquatic insects revealed by DNA barcoding

**DOI:** 10.1101/2023.11.07.565981

**Authors:** JN Negishi, MK Alam, K Tojo, F Nakamura

**Affiliations:** Faculty of Environmental Earth Science, Hokkaido University, N10 W5 Sapporo, Hokkaido 060-0810, Japan; Graduate School of Environmental Science, Hokkaido University, N10 W5 Sapporo, Hokkaido 060-0810, Japan; Biology Department, Faculty of Sciences, Shinshu University, Asahi 3-1-1, Matsumoto, Nagano 390-8621, Japan; Research Faculty of Agriculture, Hokkaido University, N9 W9 Sapporo, Hokkaido 060-8589, Japan

**Keywords:** Hyporheic zone, EPT, riparian vegetation, subsidy, insect emergence

## Abstract

How subsurface hyporheic zone (HZ) of rivers is connected to riparian zones remains largely unknown. We collected benthic macroinvertebrates and adult aquatic insects for six years, including those at 30-cm depth in the HZ to identify insect taxa having a high level of habitat affinity for HZ (HZ taxa). Adults of HZ taxa were identified with the aid of CO1 gene barcoding, and the relative abundance of HZ taxa in the riparian zone was quantified. In addition to the previously known stonefly *Alloperla ishikariana*, three species of stonefly Leuctridae and one caddisfly species of Philopotamidae were identified as HZ taxa. At the annual scale, HZ taxa accounted for approximately 38% of the total aquatic insects in the family of stoneflies (Plecoptera), caddisflies (Trichoptera), and mayflies (Ephemeroptera) in abundance and 26% of biomass, with their seasonal peaks in early spring and mid-summer (60% in abundance and 47% in biomass). Few individuals of HZ taxa were found in benthic samples (<0.1%), and hyporheic samples quantified more but erroneously estimated relative abundances of adult HZ taxa, with Leuctridae and Philopotamidae being substantially underrepresented relative to their adult abundance. Cryptic biological connectivity between subsurface and riparian zones via winged HZ-taxa adults is substantial. DNA-based species identification combined with community surveys of adult HZ-taxa complementarily used with benthic data can substantially improve the effectiveness of biomonitoring programs and outcomes of habitat conservation based on more complete picture of ecosystem health.

## Introduction

Current views of natural environments at the landscape scale stress the key roles of a mosaic of various habitat or ecosystem patches and their connectivity in maintaining populations of organisms by allowing their movements and increasing fitness (Correa Ayram et al., 2016). Another biological connection among environmentally contrasting patches is through ecosystem subsidies, defined as material fluxes such as food resources for consumers at ecosystem boundaries (Polis et al., 1997; Loreau et al., 2003). These spatial perspectives on landscape ecological properties apply to inland aquatic systems. In lakes and ponds, coastal wetlands and littoral and offshore zones are characterized by unique biological communities and resource and animal movements across them (Schoen et al., 2016; O’Reilly et al., 2023) whereas riparian zones along the shore interact with aquatic compartments through ecosystem subsidies (Gratton et al., 2008; Francis & Schindler, 2009; Jonsson & Wardle, 2009). Similarly, in rivers, hierarchically nested habitats range from microhabitats in pools and riffles to river segments within an upstream-downstream continuum (e.g., Poole, 2002; Gurnell et al., 2016). At river riparian boundaries, aquatic insects play a significant role in connecting terrestrial consumers with allochthonous food prey (Baxter et al., 2005). The presence of riparian zones is also important because aquatic insects mate and reproduce there to eventually return to the river (Petersen et al., 2004; Rahman et al., 2021).

Despite of well-established notion that rivers have a vertical dimension in the subsurface zone (Stanford & Ward, 1993), the magnitude of subsurface connectivity to riparian zones remains unclear. The functional subsurface zone in rivers is known as the hyporheic zone (HZ), where surface and groundwater mix (Boulton, 2007; Marmonier et al., 2012; Lewandowski et al., 2019). One of the HZ key ecological functions is habitat provisioning to invertebrate communities, including insect taxa, for temporary or persistent use (Stanford & Gaufin, 1974; Malard et al., 2003; Stubbington et al., 2011; Mathers et al., 2017; Dunscombe et al., 2018; Dorff & Finn, 2019; Negishi et al., 2019a). Adults leave water; thus, taxa with a high habitat affinity for HZ can serve as an indicator of biological connectivity between the subsurface and riparian zones (Alam et al., 2020). Several studies have estimated invertebrate production in the HZ at the community (Smock et al., 1992) or species levels (Collier et al., 2004; Wright-Stow et al., 2006) in relatively small, forested rivers. Seminal theoretical frameworks have advocated that the extent of the hyporheic habitat and its functional significance should be maximized in areas with sediment deposits and river braids, typically in gravel-bed rivers on alluvial plains (Stanford & Ward, 1993). Studies on the Flathead River of Montana, USA and Satsunai River of Hokkaido, Japan, valuable, both suggesting that there are potentials of high levels of resource linkages between subsurface zone and terrestrial zones (DelVecchia et al., 2016, 2019; Rahman et al., 2021a). However, quantitative year-round community-level estimates are lacking.

Exploring the structure and function of the HZ has posed a challenge because of the nature of the zone as being physically concealed under the riverbed sediment and hardship involved in efficient and frequent observations (Palmer, 1993). For instance, identifying hyporheic taxa and/or their habitat use requires intensive sampling of invertebrate communities in both the benthic zone and the HZ to assess their preference and habitat affinity levels (Stubbington et al., 2011; Negishi et al., 2019a, 2022). An increasing number of studies have taxonomically identified hyporheic insects; however, limited information exists, particularly for larvae in rivers, particularly in large floodplain rivers and their aquifers. Thus, it is imperative to improve the database of hyporheic taxa to advance our understanding of hyporheic zones, communities, and their functions (Boulton et al., 2010).

Another technical difficulty is that taxonomic keys are less established for hyporheic organisms because larvae can be caught less efficiently. In addition, some individuals spend an early life state in the HZ, which makes morphological identification problematic (Malard et al., 2003). In light of the challenges associated with the scarcity of morphological identification keys, recent developments in techniques using genetic barcoding markers can substantially facilitate species identification, including adult-larva relationships of aquatic insects and meiofaunal communities (Ruiter et al., 2013; Weigand et al., 2018; Chen et al., 2023) regardless of their size and incomplete larval taxonomic keys. Although this field has great potential, there have been limited cases of hyporheic insects (Torii et al., 2022).

The objective of this study was to quantify the level of biological connectivity between the HZ and riparian zones via winged aquatic insects in order to fill the knowledge gap in our spatial understanding of subsurface zone roles in river-riparian resource transfer and freshwater invertebrate community ecology. We focused on flying insects in the riparian zone for two reasons: First, the importance of river-riparian connectivity has been emphasized because of the animal movements between them (Baxter et al., 2005; Richardson et al., 2010). Second, previous studies have suggested a relatively high abundance of hyporheic insects in the riparian zone relative to open water away from the riparian zone (Rahman et al., 2021a). We first identified species with a high level of habitat affinity for the HZ (hyporheic species) based on aquatic larval data. In addition to the previously identified hyporheic stonefly species, *Alloperla ishikariana* (Order: Chloroperlidae) (Negishi et al., 2019a, 2022), we sought to identify several more species to increase the level of credibility of our estimates by referring to benthic invertebrate data in different seasons over five years. We then collected adult insects and estimated them throughout a year over six years to understand the contribution of hyporheic species to the total abundance and biomass in the riparian zone. Six-year monitoring data were used to represent reliable average estimates, because insect emergence fluctuates among and within years (Negishi et al., 2022). DNA-based barcoding analyses were used to conclusively relate larvae to adults and their taxonomic identifications. We predicted that hyporheic insects are relatively more abundant in the riparian-vegetation zone than the estimates inferred from benthic invertebrates.

## Materials and methods

### Sites

This study was conducted from June 2015 to October 2022 in the Satsunai River (catchment area, 725 km^2^), which runs from Mt. Satsunai (42^°^41 N, 142^°^47 E; 1,895 m above sea level) to the Tokachi River, Eastern Hokkaido, Japan (Supplementary information 1). In total, 11 sites (50–500 m long channel reaches) were chosen (10 sites in the main channel and one site in a tributary stream) within a 15-km section to capture diverse invertebrate communities and taxa in the area. There is a gradient in water quality from upstream to downstream due to the presence of a waste water treatment plant and non-point sources of nutrients in surrounding agricultural fields (Negishi et al., 2019a, b.). This study used data irrespective of such environmental variabilities so that average estimates of significance of hyporheic insects in flying adults insects in riparian zone in the area can be obtained. The regional climate was characterized by the lowest air temperature and precipitation in winter and the highest temperatures and precipitation in summer to autumn. The mean (±standard deviation) annual precipitation and temperature from 2009–2019 were 1,207.5 (±202.2) mm and 5.5 (±0.3) °C, at Kami-Satsunai observation station (Japan Meteorological Agency; 200 m away from the study reach). The mean (±SD) monthly air temperature for the winter months (December, January, and February) and the summer months (June, July, and August) for the same period at the same station were -7.1 °C (±2.0) and 16.9 °C (±2.5), respectively. More detailed site information can be found in previous studies (Negishi et al., 2019a,b; Alam et al., 2020, 2021).

### Sample collection

We used the invertebrate data collected by the Ministry of Land, Infrastructure, Transport, and Tourism (MLIT) as a part of environmental river assessments in relation to the Satsunai-gawa dam (Nakamura et al., 2020). Benthic invertebrate communities were collected at two sites (G and H) to characterize their community structure of benthic invertebrates, as described by Negishi et al. (2019b). Samples were collected in three seasons in two sections (each approximately 200-300 m), each of which corresponded to habitat type (main or secondary channels), at each of the two sites over five years, and some seasonal and site collections were not conducted due to funding limitations (Supplementary information 2A). Three replicates of benthic invertebrate samples were collected using Surber samplers (25 cm × 25 cm; 475-µm mesh) at randomly selected locations within a representative riffle of each section, providing a total of 114 samples.

Hyporheic macroinvertebrates were collected using hyporheic traps in the glide habitat at four sites (S, A1, L1, and L7) to identify the hyporheic species. Sampling reaches (30-50 m) were set at the upstream and downstream ends of sites, with a between-reach distance ranging from 100 m at site S to 2.86 km at site A1. Each reach was subdivided longitudinally into two sub-reaches, 10 m apart. At each sub-reach, one pit was excavated to a depth of 30 cm using either an excavator or a hand. Two traps were installed at each pit. Trap installations were conducted in August 2020 and were manually excavated and retrieved using a hand-held D-frame net (375-μm mesh) in three months since the installation (Supplementary information 2 B). The traps were retrieved after removing sediment from the trap. Details of the trapping methods can be found in Alam et al. (2021). All samples at the downstream reach of site A1 were lost due to floods and were excluded from further analyses (28 hyporheic samples). Immediately before retrieving traps, the riverbed surface of each pit above the buried traps was sampled using a Surber sampler, as described above in the MLIT survey (a total of 14).

Adult aquatic insects were caught using single-headed Malaise traps (length:165 cm, width:180 cm, height:180 cm; Megaview Science Co., Taichung, Taiwan) set at six sites along the edge of rivers between March and October in six years (2017–2022) as previously conducted by Rahman et al. (2021a) and Negishi et al. (2022). The traps were arranged such that the long axis was parallel to the river channel and the distance to the water edge was within 3 m, and the 75%ethanol preservation bottles were replaced at 3-48 day intervals (Supplementary information 2C). During non-trapping periods, especially in the winter months (November to early January), we conducted visual surveys of the riparian zone and river-channel edges to ensure that the target groups did not emerge or fly. However, from late January to March, stoneflies in Capniidae emerged and were not fully measured because malaise traps could not be used in snowing weather conditions. In some years, sampling was not conducted throughout the period owing to logistical limitations, providing a total of 366 samples. Emergence traps set on water surface were not used because the several-year application of such traps in the study river system, where frequent changes in river channel morphology and water prevail, was impossible.

### Invertebrate processing

All invertebrate larvae collected were rinsed from the substrate materials and preserved *in situ* either using a 10% formaldehyde solution after triplicate samples were pooled together (MLIT samples) or using 75% ethanol without sample pooling (other data). Non-MLIT samples were sieved through a 500-µm mesh before preservation, whereas MLIT samples were larger than the 475-µm, the size of sampler’s mesh size. Upon returning to the laboratory, invertebrates were sorted under a stereoscopic microscope, enumerated, and identified to the lowest level possible using various taxonomic keys (e.g., Maruyama & Takai, 2000; Kawai & Tanida 2005). Adult invertebrates in collection bottles were brought back to the laboratory and sorted and identified to species, family, or order levels, as done for other samples. For genetic analyses, some individuals were transferred to 99.5% ethanol and stored until further analyses.

### DNA barcoding

The larvae of taxa with a high affinity for the hyporheic zone were examined for their adult-larvae associations based on DNA barcoding using DNA sequences in the CO1 gene region. Non-MLIT larval and adult samples and previously collected samples were used for this purpose, and the taxonomic identity of the sample was determined based on morphological keys. Tissues of hindleg muscles were removed from 99.5%-ethanol preserved individuals and their total genomic DNA was extracted and purified using the DNeasy-Tissue-Kit (QIAGEN, Hilden) or individually purchased reagents, including cell lysis solution, proteinase *k*, and protein precipitation solution. The extracted DNA was then amplified for the 658-bp CO1 region fragments by polymerase chain reaction (PCR) with universal primers LCO1490 and HCO2198 (Folmer et al., 1994) (Supplementary information 3). The PCR products were purified using ExoSAP-IT Express (Thermo Fisher Scientific Japan, Tokyo) or illustra ExoProStar (GE Healthcare, Amersham, UK), or individually purchased reagents using a PEG-based purification procedure. Purified DNA fragments for the mtDNA COI region were sequenced on an Applied Biosystems 3130xl instrument using the BigDye Terminator v3.1 Cycle Sequence Kit (Applied Biosystems, Waltham, MA USA).

### Analyses

We calculated Manly’s selectivity ratio for each habitat type (benthic or hyporheic) using the hyporheic trap study as the habitat affinity index (HAI) of the EPT taxa (Heisey, 1985). This index considers the availability of each habitat type in the environment. Abundance data were converted to presence or absence data for each sample and the ratio was adjusted for uneven habitat-type availability. A value greater than 0.5 indicated a positive selectivity of hyporheic habitat over benthic habitat.

The own DNA sequences were combined with those of closely related species in the GenBank database (NCBI). Closely related species sequences were found in the BLAST search, and those belonging to insects with >90% identity to the searched taxa were downloaded. These sequences were aligned and truncated at maximum common base-pair length (596 bp) and used to construct a phylogenetic tree based on the neighbor-joining (NJ) method (Saitou & Nei, 1987) with a rapid bootstrapping procedure using 1,000 bootstrap analyses using MEGA X (Kumar et al., 2018). Based on the adult community data, the annual pattern of emergence was statistically obtained using generalized additive models (GAMs) at appropriate taxonomic resolutions. As an explanatory variable, Julian days (middle day for respective sampling durations) were used, and the response variables were abundance and biomass per day per trap, respectively. Biomass was obtained by multiplying the abundance data with taxon-average dry weight (g) data (previously reported in Rahman et al., 2021a, and partially unpublished data of JNN). Based on the habitat-type affinity, the abundance or biomass was summed for hyporheic and benthic species, and the proportion of hyporheic species was obtained in each sample; these data were also analyzed by GAMs.

All analyses were conducted using R (R Core Team, 2022) and relevant packages, such as *ggtree*, *ggplot2*, and *mgcv* (Wickham & Wickham, 2016; Wood, 2017; Yu et al., 2017).

## Results

We caught 21,117 adults, 102,526 larvae in the MLIT data, and 6,315 larvae in benthic and hyporheic samples (Supplementary information 4, 5, 6). Three taxa, Philopotamidae, Molannidae, and Leuctridae, were found only in HZ (HAI=1), with Polycentropodidae and *A. ishikariana* demonstrating high to marginally high hyporheic-habitat affinity (Fig. 1). Philopotamidae, Leuctridae, and *A. ishikariana* were abundant in adults, comprising 11–21% of the total catch. Philopotamidae adults were morphologically identified as belonging to the genus *Kisaura* whereas leuctrid adults comprised three species (*Paraleuctra cercia*, *Perlomyia* sp.1, and sp.2). Benthic invertebrates were dominated by grazers and collector-gatherer mayflies (e.g., Heptegeniidae) and filter-feeder caddisflies (e.g., Stenopsychidae). Based on the disproportionately high abundance in the riparian zone with high HAI, three taxa (Philopotamidae, Leuctridae, and *A. ishikariana*) were chosen as focal HZ species.

**Figure 1.**
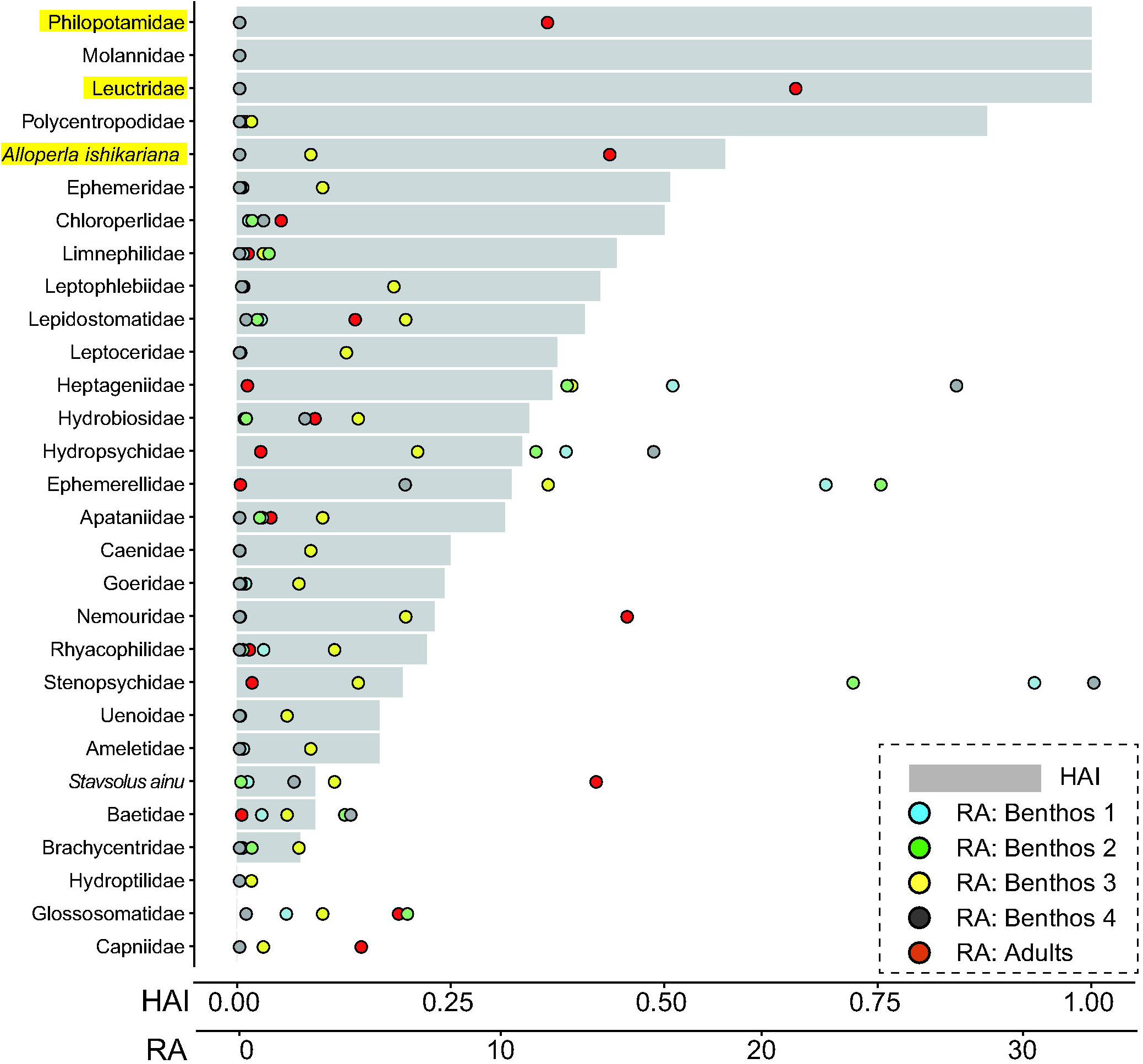
EPT taxa with their habitat affinity index (HAI) based on Manly’s selectivity ratio and their relative abundance (RA) in four larval and one adult EPT dataset. Benthos 1: MLIT data in June; Benthos 2: MLIT data in July; Benthos 3: non-MLIT data in November; and Benthos 4: MLIT data in October. Chloroperlidae includes individuals other than *Alloperla ishikariana*. Three taxa highlighted in yellow were the focal hyporheic species.

The occurrence of Philopotamidae was rarer than that of the other two taxa, whereas both Philopotamidae and Leuctridae were much less abundant than *A. ishikariana* (Fig. 2). The larval Leuctridae were not morphologically distinguishable at the species level. Based on DNA barcoding using 136 sequences (Supplementary information 7), three hyporheic taxa were associated with their larvae (Fig. 3). All larvae identified as *A. ishikariana* were included in a distinct clade with a long branch length together with their adults. Leuctrid larvae sequences were assigned to three clades, corresponding to the three adult species. Likewise, both adults and larvae of Philopotamidae were included in the same clade that deeply branched off from the one for closely related *Kisaura aurascens*, suggesting their adult-larva associations. Thus, HAI-based categorization of adults is reliable.

**Figure 2.**
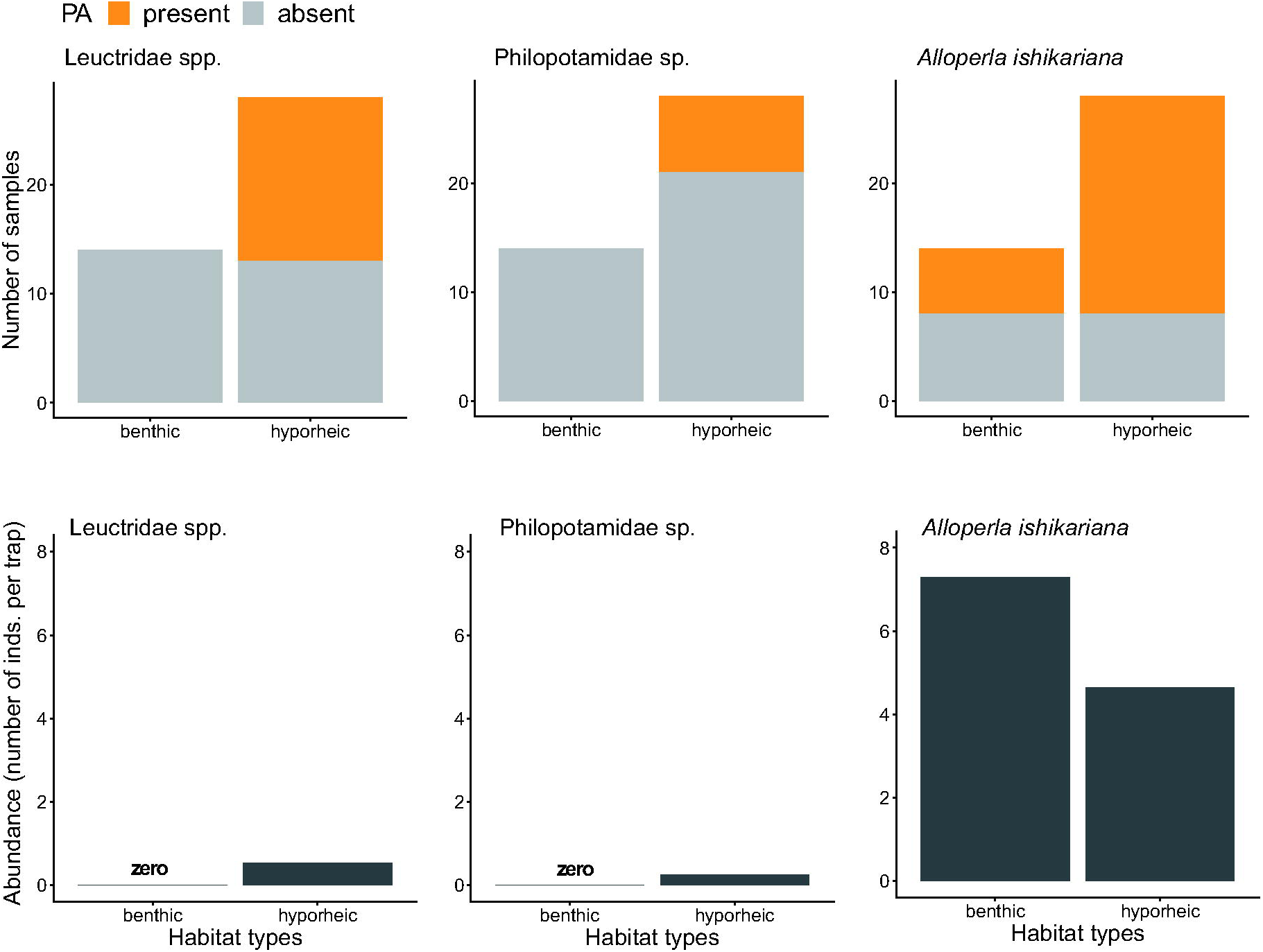
The occurrence (presence or absence, upper panels) and abundance obtained from a total individual number and total sample size (lower panels) of three taxa in hyporheic samples and simultaneously collected non-MLIT benthic samples (upper panels).

**Figure 3.**
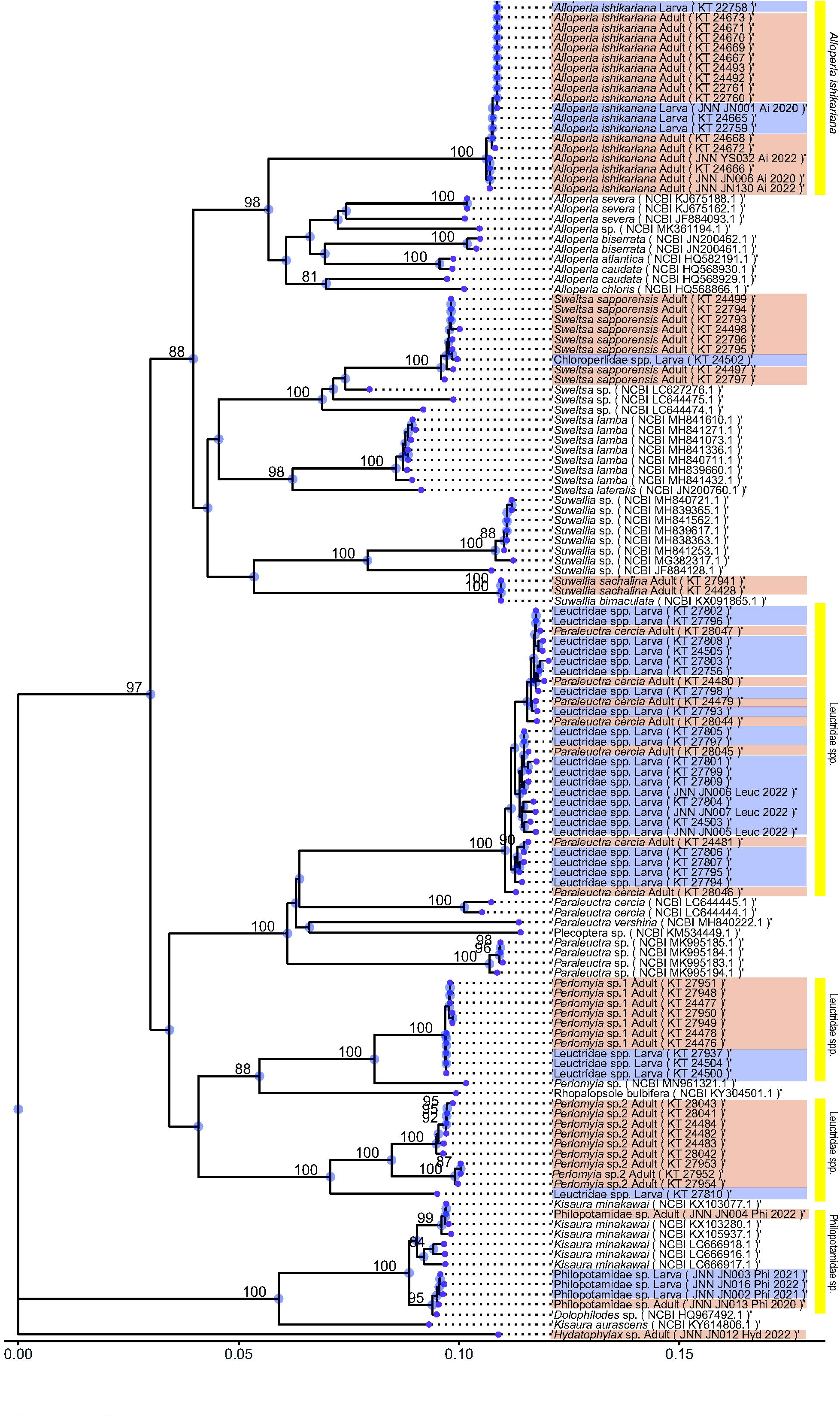
The estimated phylogenetic relationships (NJ tree) of focal taxa, Philpotamidae, Leuctridae, and *A. ishikariana* and their closely related species based on mtDNA CO1 region. The numbers at major nodes indicate bootstrap probabilities (only those >80 was shown). The scale bar indicates branch lengths in nucleotide substitutions per site. Labels shown in light blue and light orange correspond with sequences for larvae and adults. Yellow bands indicate the range of branch ends for hyporheic taxa considered in adult EPT category for their habitat affinity. The sequences labeled with NCBI were obtained from GenBank database; those labeled with JNN and with KT were sequenced at Hokkaido University and Shinshu University, respectively; “Larva” and “Adult” denote life-cycle stage of samples. Hydatophylax sp. Adult (JNN JN012_Hyd_2022) was added to form a root outgroup of the tree.

Most of the numerically abundant taxa were significantly modelled by Julian days in GAMs (Table 1). The peak of Leuctridae occurred in early May, and two other species emerged in mid-summer (Fig. 4). Two peaks, a sustained peak in early June and another in early July, were observed for *A. ishikariana*, whereas the other two showed unimodal emergence peaks. A variety of peak patterns were observed for the other benthic taxa. The mean abundance proportion associated with HZ to total abundance was 37.9%, with a peak of 59.8%. The mean biomass proportion of hyporheic taxa was 26.5%, with a peak of 47.7%, because some benthic taxa such as Perlodidae (*Stavsolus ainu*) had large per-capita biomass. Two peaks in the proportion of hyporheic taxa were apparent in mid-May and early July.

**Figure 4.**
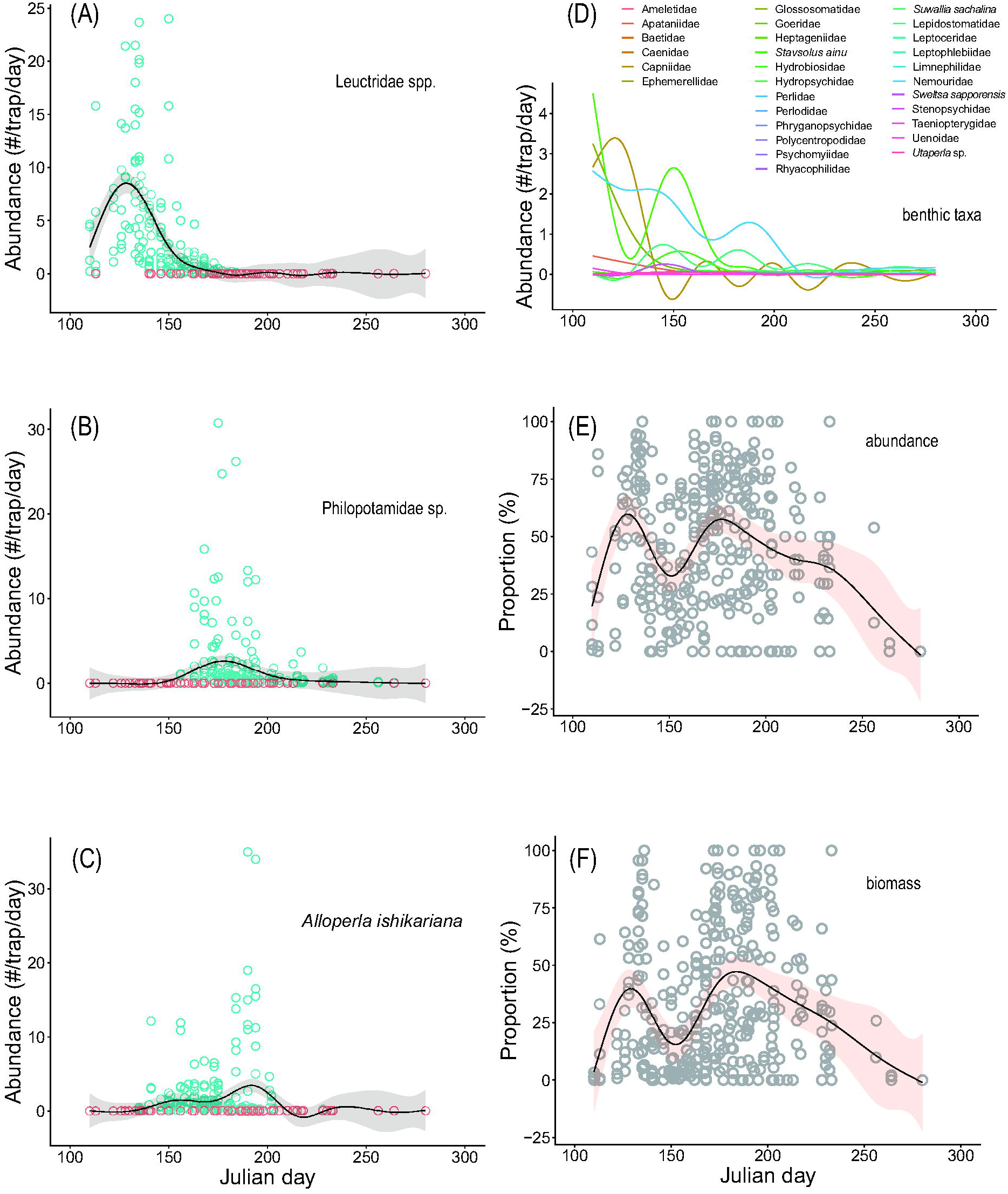
Emergence patterns of adults of three hyporheic taxa in relation to Julian day (A, B and C). Regression lines based on generalized additive models (GAMs) were shown in solid lines with 95% confidence intervals shown as gray clouds; blue and red open circles represent data with and without the catch of individuals. In (D), the model regression lines of GAMs for other non-hyporheic benthic taxa were shown. The proportions of hyporheic taxa compared to non-hyporheic benthic taxa were shown for abundance (E) and biomass (F). Regression lines based on GAMs were shown in solid lines with 95% confidence intervals shown as light orange clouds.

**Table 1.** Results of generalized additive models explaining the hourly catch of adult aquatic insect, and proportions of hyporheic species in total abundance and biomass (%abundance and %biomass) in relation to Julian day. For each response variable, F-values and statistical significance of explanatory variable (p), generalized cross-validation index (GCV), and percentage of total deviance explained (%) were shown. No asterisk indicates p>0.05, and the number of asterisks corresponds to model significance; *p<0.05, **p<0.01, and ***p<0.001.

| Response variables | F-values | p | GCV | Explained deviance (%) |
| --- | --- | --- | --- | --- |
| Leuctridae | 36.64 | *** | 8.90092 | 0.48 |
| Philopotamidae | 5.52 | *** | 10.26288 | 0.11 |
| *Alloperla ishikariana* | 4.32 | *** | 12.75260 | 0.11 |
| Ameletidae | 1.07 |  | 0.00068 | 0.02 |
| Apataniidae | 18.93 | *** | 0.05110 | 0.20 |
| Baetidae | 3.00 |  | 0.00148 | 0.01 |
| Caenidae | 0.34 |  | 0.00045 | 0.00 |
| Capniidae | 8.17 | *** | 4.43198 | 0.17 |
| Ephemerellidae | 1.93 |  | 0.00071 | 0.01 |
| Glossosomatidae | 13.15 | *** | 1.34839 | 0.22 |
| Goeridae | 0.99 |  | 0.00263 | 0.02 |
| Heptageniidae | 3.17 | ** | 0.00561 | 0.08 |
| *Stavsolus ainu* | 11.46 | *** | 3.72775 | 0.23 |
| Hydrobiosidae | 5.92 | *** | 0.28254 | 0.13 |
| Hydropsychidae | 2.57 | * | 0.02806 | 0.06 |
| *Sweltsa sapporensis* | 6.08 | *** | 0.00810 | 0.13 |
| Lepidostomatidae | 3.05 | ** | 0.71326 | 0.07 |
| Leptoceridae | 1.73 |  | 0.00000 | 0.04 |
| Leptophlebiidae | 2.35 | * | 0.00315 | 0.04 |
| Limnephilidae | 3.49 |  | 0.00428 | 0.08 |
| Nemouridae | 5.39 | *** | 4.25207 | 0.11 |
| Perlidae | 3.83 | ** | 0.00098 | 0.05 |
| Perlodidae | 1.56 |  | 0.00003 | 0.00 |
| Phryganopsychidae | 8.63 | *** | 0.00001 | 0.18 |
| Polycentropodidae | 5.24 | *** | 0.00250 | 0.07 |
| Psychomyiidae | 1.73 |  | 0.00002 | 0.04 |
| Rhyacophilidae | 4.05 | ** | 0.01082 | 0.06 |
| *Suwallia sachalina* | 8.03 | *** | 0.03833 | 0.17 |
| Stenopsychidae | 2.89 | ** | 0.01545 | 0.06 |
| Taeniopterygidae | 15.47 | *** | 0.00153 | 0.28 |
| Uenoiidae | 0.23 |  | 0.00043 | 0.00 |
| *Utaperla* sp. | 2.45 | * | 0.01774 | 0.03 |
| %abundance | 8.35 | *** | 700.92090 | 0.18 |
| %biomass | 8.46 | *** | 756.09260 | 0.18 |

## Discussion

Our prediction was supported because hyporheic adult insects were among the most numerically abundant in the riparian vegetation zone, in sharp contrast to the exceedingly low abundance found in benthic invertebrate data. Importantly, benthic invertebrate data that has been used as baseline information of river habitat monitoring provided completely different faunal picture compared with that in our Malaise trap data. Results based on field observations over more than five years provide significant management and conservation implications, as well as scientifically advanced understanding of ecological processes in lowland gravel-bed rivers.

Strong HZ affinity was demonstrated for Leuctridae and Philopotamidae. Leuctrid species are known to be hyporheic dwellers (Weigelhofer & Waringer, 2003; Dorff & Finn, 2020; Malison et al., 2020; Negishi et al., 2022). DNA-barcoding conclusively demonstrated that leuctrids comprise three species. Lrvae of *Paraleuctra cercia* and *Perlomyia* sp.1 were identified and assigned to the clade of their adults. The sequence of one larva (KT27810) was found to be slightly divergent from the clade of *Perlomyia* sp.2. Other adults identified as *Perlomyia* sp.2 also showed some differences. No other morphologically distinct adults were found in the leuctrids. Therefore, the larva (KT27810) was assigned as *Perlomyia* sp.2. This species was relatively minor in Leuctridae (in adults, 8.7% of the total number of Leuctridae) with a relatively small body (smaller by >50% than the two other species); thus, the bias of hyporheic insect abundance and biomass caused by the inclusion of this species was minor. It was only recently that elusive larvae of Philopotamidae (*Kisaura minakawai*) were found in the HZ (Torii et al., 2022). Likewise, our extensive field sampling found only a few larvae of Philopotamidae in the HZ at a very low abundance. DNA barcoding suggested that our specimen belonged to the *Kisaura* genus, likely *K. minakawai*. More *Perlomyia* sp.2 and Philopotamidae larvae need to be collected to enhance the DNA database for key morphological developments. Larvae of *A. ishikariana* have been largely found in the HZ (Negishi et al., 2019a; 2022), which is a discrepancy with our observations. In this study, many of them on the surface were first instars not very long since their hatching, whereas those found in HZ were mixtures of first instars and larger individuals in older cohorts, which suggests that we identified the approximate timing of newly hatched instars immediately before their migration into HZ (Negishi et al., 2022).

Our previous study in the same areas provided the only available matter flux estimate from HZ to riparian zone based on carbon and nitrogen in single species *A. ishikariana* adults (Rahman et al., 2021a), and reported the HZ-originated potential resource for consumers as high as 70% at their peak emergence in the summer. Although our community-level results did not quantify resource transfer, they suggest that the abundance of HZ-derived adult aquatic insects and resource contribution from the hyporheic zone to the riparian zone can be potentially high even in early spring. This is important because the total amount of resources from rivers can be at its highest in early spring (Rahman et al., 2021), and terrestrial resources are relatively low in spring (Nakano & Murakami, 2001). How to quantify the amount of HZ resources assimilated into the food web is a critical future research question. Studies on the Nyack River are relevant and implicative in advancing this methodology. Methane-derived carbon resources were assimilated by aquatic consumers inhabiting shallow aquifers below large floodplains, indicating the importance of subsurface productivity (DelVecchia et al., 2016, 2019). This was possible because methane-based carbon resources were labeled with characteristically depleted carbon isotope ratios, and the fate of this resource could be theoretically tracked by measuring the isotope contribution to land animals and plants. However, this method may be applicable to a limited number of cases because unique isotopic signatures are not necessarily common as seen in *A. ishikariana* (Alam et al., 2020).

Abundant hyporheic species in the riparian zone point to the importance of the riparian zone for their population dynamics through dispersal and habitat provisioning to mating individuals. For example, *A. ishikariana* preferentially moves into the riparian zone in the lateral direction and then longitudinally over several kilometers (Rahman et al., 2021a, b), suggesting that they depend on the riparian zone as their dispersal corridor (see also Macneale et al., 2005). This is consistent with the notion that riparian zone usage and thus conditions of the riparian zone are potentially important in the organization of river invertebrate communities (Delettre & Morvan, 2000; Arce et al., 2023; Palt et al., 2023). Some of these previous studies examined benthic invertebrates in relation to riparian vegetation structure (Arce et al., 2023; Palt et al., 2023). Our findings highlight the potential risk of underestimating the associations between riverine invertebrates and riparian zone structures if hyporheic species are not properly identified. As clearly shown in this study, hyporheic taxa can be overlooked if they are based only on benthic larval samples. The importance of riparian vegetation structure and connectivity for flying aquatic insects is expected to be greater in lowland rivers where human activities intensively fragment landscapes (Palt et al., 2023). Furthermore, because contemporary bio-assessment programs are largely based on benthic taxa (those largely found on the surface zone of riverbed) (e.g., Haase et al., 2004), the environmental conditions cannot be associated in terms of hyporheic environment, leading to erroneous status indications of river ecosystems in biomonitoring programs and/or target-setting in restoration projects (Boulton et al., 2010; Alam et al., 2021). Communities in hyporheic and benthic zones tend to respond differently to environmental stressors or restoration activities (Negishi et al., 2019a; Robertson et al., 2021).

Therefore, future research in the field should examine the relationships between river and landscape characteristics and community structure, including hyporheic communities. Data of conventional benthic samples can be complementarily integrated with data of flying adults to substantially advance the understanding of river ecosystems.

Leuctidae and Philopotamidae were extremely rare in hyporheic traps, despite their popularity in the adult community. In addition, Capniidae were hardly caught in benthic samples while their adults were relatively abundant. The following central question arises. How and where do these adults emerge? Hyporheic communities differ among depths and horizontal spatial extents (Malard et al., 2003; Weigelhofer & Waringer, 2003; Davy-Bowker et al., 2006), and our sampling of hyporheic communities only in glides at a fixed depth near the water edge is the key reason behind the above-mentioned discrepancy. Currently, we do not have data to identify their likely source habitats, and can only propose potential source habitats. First, areas that are laterally away from the main channel are possible. Stoneflies emerge from an area several 100 m away from the main channel of Flathead River in the Nyack floodplain system (DelVecchia et al., 2016, 2019). We did not have extensive underground monitoring well systems extending to that far from the river channel. Another possibility is a hydrologically active zone near riffles. Upwelling and downwelling areas have different community structures (Davy-Bowker et al., 2006). In addition to being fixed at a single depth, our method cannot evaluate the depth-integrated habitat availability of the HZ, which provides a three-dimensional habitat for animals. Benthic monitoring almost completely missed them and highlighted the pitfall of underestimating secondary production as well as species diversity. These unsampled areas are difficult to examine physically; installations and maintenance of monitoring well to cover a wide range of depth as well as distant areas from the channel would be challenging. Thus, new technologies such as eDNA of surface and subsurface water might have potential.

In conclusion, we demonstrated cryptic biological connectivity between the subsurface and riparian zones through the winged adults of hyporheic taxa. Our findings highlight the potential pitfalls in the ecological understanding of gravel-bed river systems with well-functioning HZ, if assessed only based on benthic samples. Pitfalls are generated because adult hyporheic insects are abundant in the riparian zone, and benthic invertebrate monitoring rarely detects their larvae.

Missing this hyporheic-riparian biological connectivity can lead to mismanagement of both the riparian zone and the HZ, as key processes driving biodiversity, material cycling, food webs, and animal dispersals associated with hyporheic insects remain cryptic and could be unknowingly degraded. Therefore, we argue that the conservation values of the HZ extend beyond the riparian-river boundaries. DNA-based species identification combined with community surveys of adult HZ-taxa can substantially improve the knowledge of the structure and function of rivers, the effectiveness of biomonitoring programs, and outcomes of habitat conservation and protection.

Although the results were limited to a single river, the Satsunai River is one of the rivers where high-quality subsurface habitats have been preserved, and thus providing a reference river system with which other river systems can be compared.

## Acknowledgements

We are grateful to the Obihiro Regional Office of Hokkaido Development Bureau, MLIT for data collection and field assistance. This study was partly supported by the research fund for the Tokachi and Ishikari rivers provided by MLIT (18056588) and JSPS KAKENHI (18H03408 and 18H03407).

## Ethical approval

No ethical violation was occurred in this research.

## Conflict of interest

Authors declare that there is no conflict of interest to disclose.

## Data availability statement

All the data used in the paper will be deposited in Dryad and DNA data bank of Japan.

## Author Contribution Statement

JN Negishi: Conceptualization (equal), data collection (equal), data curation (equal), formal analysis (equal), funding acquisition (equal), and leading writing. MK Alam: Data collection (equal), data curation (equal), and editing draft manuscript (equal). K-Tojo: Conceptualization (equal), data curation (equal), formal analysis (equal), editing draft manuscript (equal), and funding acquisition (equal). F Nakamura: Conceptualization (equal), editing draft manuscript (equal), and funding acquisition (equal). All the authors contributed critically to the drafts and approved the final manuscript for publication.

## Notes

### Competing Interest Statement

The authors have declared no competing interest.

