## Supplementary material for "Cryptic connectivity between hyporheic and riparian zones via winged aquatic insects revealed by DNA barcoding": SI1

**Supplementary information 1**

Study site map for field sampling. The location of study area were shown in a dotted square in relation to Tokachi River watershed in Hokkaido; blue lines denote rivers whereas the white to black gradation indicate elevation above sea level (A). The location of study sites in Satsunai River; a star indicates Satsunai-gawa dam (B).

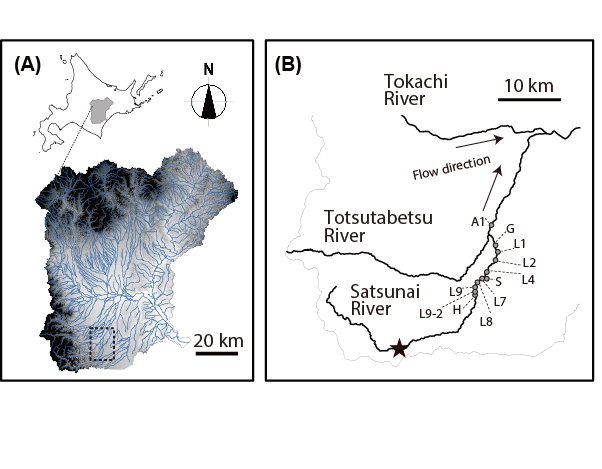
