## Supplementary material for "Cryptic connectivity between hyporheic and riparian zones via winged aquatic insects revealed by DNA barcoding": SI2

**Supplementary information 2A**

Summary of benthic sample collections of the MLIT monitoring survey in relation to season, year, and habitat. NA denotes that samples were not collected due to funding limitations. The main channel section was the largest water course, whereas the secondary-channel section was the second largest one that branched off from the main channel. The former was deeper with higher average current velocity relative to the latter.

| Site | Habitat | 2015 | 2016 | 2017 | 2018 | 2019 |
| --- | --- | --- | --- | --- | --- | --- |
| G | Main channel | June, July, October | June, July | June, July | June, July | June, July |
|  | Secondary channel | June, July, October | June, July | June, July | June, July | June, July |
| H | Main channel | NA | June, July | June, July | June, July | June, July |
|  | Secondary channel | NA | June, July | June, July | June, July | June, July |

**Supplementary information 2B**

Summary of sample collection of benthic and hyporheic samples in 2020. Down and up denote sub-reach samples collected at downstream and upstream sub-reaches, respectively.

| Site | Installation of hyporheic traps | Collection of benthic samples and hyporheic traps |
| --- | --- | --- |
| A | August 4th, 2020 | October 28th, 2020 |
| S | August 4th, 2020 | October 28th (down) and 29th (up), 2020 |
| L1 | August 4th, 2020 | October 28th, 2020 |
| L7 | August 4th, 2020 | October 29th, 2020 |

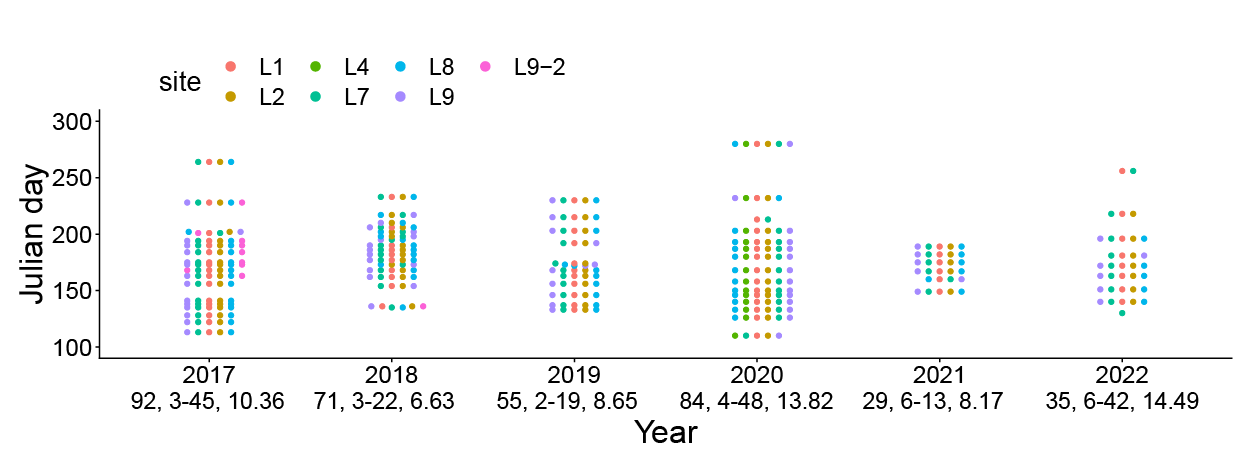
**Supplementary information 2C**

Sampling occasions of adult samples using Malaise traps, shown in points (median Julian days of sampling periods) in each year. Sample size, and range and mean of trapping intervals were shown for each year. The interval was adjusted so that frequent sampling was performed intensively in the period high insect activities.
