## Supplementary material for "Cryptic connectivity between hyporheic and riparian zones via winged aquatic insects revealed by DNA barcoding": SI3

**Supplementary information 3**

PCR conditions in the present study. Samples processed in Shinshu University (A) and samples processed in Hokkaido University (B). For the information regarding sample-PCR condition relations, please refer to the Supplementary information 7**.**

(A)

| Steps | Temperature (℃) | Duration (min) | Cycle repeat | Cycle times |
| --- | --- | --- | --- | --- |
| Pre-heating | 94 | 5 |  |  |
| Denaturing | 94 | 1 | Yes | x35 cycles |
| Annealing | 45 | 2 | Yes |  |
| Extending | 72 | 1 | Yes |  |
| Final extending | 72 | 7 |  |  |
| Storing | 4 | ∞ |  |  |

(B)

| Steps | Temperature (℃) | Duration (min) | Cycle repeat | Cycle times |
| --- | --- | --- | --- | --- |
| Pre-heating | 95 | 2 |  |  |
| Denaturing | 94 | 1 | Yes | x 25 cycles |
| Annealing | 45.5 | 0.5 | Yes |  |
| Extending | 72 | 1 | Yes |  |
| Final extending | 72 | 5 |  |  |
| Storing | 10 | ∞ |  |  |
