## Supplementary material for "Cryptic connectivity between hyporheic and riparian zones via winged aquatic insects revealed by DNA barcoding": SI4

**Supplementary information 4**

Total abundance and relative abundance (%) of adult EPT individuals caught in this study

| Order | Family | Species | Abundance | Relative abundance |
| --- | --- | --- | --- | --- |
| Ephemeroptera | Heptageniidae |  | 63 | 0.30 |
| Ephemeroptera | Leptophlebiidae |  | 26 | 0.12 |
| Ephemeroptera | Baetidae |  | 17 | 0.08 |
| Ephemeroptera | Ameletidae |  | 6 | 0.03 |
| Ephemeroptera | Ephemerellidae |  | 6 | 0.03 |
| Ephemeroptera | Caenidae |  | 3 | 0.01 |
| Plecoptera | Leuctridae |  | 4498 | 21.30 |
| Plecoptera | Nemouridae |  | 3135 | 14.85 |
| Plecoptera | Chloroperidae | *Alloperla ishikariana* | 2994 | 14.18 |
| Plecoptera | Perlodidae | *Stavsolus ainu* | 2883 | 13.65 |
| Plecoptera | Capniidae |  | 984 | 4.66 |
| Plecoptera | Chloroperidae | *Suwallia sachalina* | 141 | 0.67 |
| Plecoptera | Chloroperidae | *Sweltsa sapporensis* | 114 | 0.54 |
| Plecoptera | Chloroperidae | *Utaperla* sp. | 81 | 0.38 |
| Plecoptera | Taeniopterygidae |  | 32 | 0.15 |
| Plecoptera | Perlidae |  | 31 | 0.15 |
| Plecoptera | Perlodidae |  | 1 | 0.00 |
| Trichoptera | Philopotamidae |  | 2490 | 11.79 |
| Trichoptera | Glossosomatidae |  | 1286 | 6.09 |
| Trichoptera | Lepidostomatidae |  | 935 | 4.43 |
| Trichoptera | Hydrobiosidae |  | 610 | 2.89 |
| Trichoptera | Apataniidae |  | 253 | 1.20 |
| Trichoptera | Hydropsychidae |  | 171 | 0.81 |
| Trichoptera | Stenopsychidae |  | 101 | 0.48 |
| Trichoptera | Rhyacophilidae |  | 79 | 0.37 |
| Trichoptera | Limnephilidae |  | 69 | 0.33 |
| Trichoptera | Polycentropodidae |  | 47 | 0.22 |
| Trichoptera | Goeridae |  | 44 | 0.21 |
| Trichoptera | Phryganopsychidae |  | 6 | 0.03 |
| Trichoptera | Uenoiidae |  | 6 | 0.03 |
| Trichoptera | Psychomyiidae |  | 4 | 0.02 |
| Trichoptera | Leptoceridae |  | 1 | 0.00 |
