## Supplementary material for "Cryptic connectivity between hyporheic and riparian zones via winged aquatic insects revealed by DNA barcoding": SI5

**Supplementary information 5**

Total abundance and relative abundance (%) of larval EPT individuals caught in MLIT data in this study

| Order | Family | Abundance | Relative abundance |
| --- | --- | --- | --- |
| Ephemeroptera | Ephemerellidae | 26561 | 25.91 |
| Ephemeroptera | Heptageniidae | 17586 | 17.15 |
| Ephemeroptera | Baetidae | 2331 | 2.27 |
| Ephemeroptera | Leptophlebiidae | 156 | 0.15 |
| Ephemeroptera | Ameletidae | 106 | 0.10 |
| Ephemeroptera | Ephemeridae | 101 | 0.10 |
| Ephemeroptera | Caenidae | 2 | 0.00 |
| Plecoptera | Chloroperlidae | 462 | 0.45 |
| Plecoptera | Perlodidae | 276 | 0.27 |
| Plecoptera | Perlidae | 25 | 0.02 |
| Plecoptera | Nemouridae | 24 | 0.02 |
| Plecoptera | Leuctridae | 3 | 0.00 |
| Trichoptera | Stenopsychidae | 32257 | 31.46 |
| Trichoptera | Hydropsychidae | 13978 | 13.63 |
| Trichoptera | Glossosomatidae | 3956 | 3.86 |
| Trichoptera | Apatanidae | 954 | 0.93 |
| Trichoptera | Lepidostomatidae | 886 | 0.86 |
| Trichoptera | Uenoidae | 777 | 0.76 |
| Trichoptera | Rhyacophilidae | 723 | 0.71 |
| Trichoptera | Limnephilidae | 566 | 0.55 |
| Trichoptera | Brachycentridae | 275 | 0.27 |
| Trichoptera | Hydrobiosidae | 267 | 0.26 |
| Trichoptera | Goeridae | 191 | 0.19 |
| Trichoptera | Leptoceridae | 36 | 0.04 |
| Trichoptera | Polycentropodidae | 27 | 0.03 |
