## Supplementary material for "Cryptic connectivity between hyporheic and riparian zones via winged aquatic insects revealed by DNA barcoding": SI6

**Supplementary information 6**

Total abundance and relative abundance (%) of larval EPT individuals caught in non-MLIT data using hyporheic traps and simultaneously collected Surber samples in this study. TA: total abundance; RA: relative abundance

| Order | Family | Species | Benthic TA | Hyporheic TA | TA | RA |
| --- | --- | --- | --- | --- | --- | --- |
| Ephemeroptera | Leptophlebiidae |  | 321 | 853 | 1174 | 18.59 |
| Ephemeroptera | Heptageniidae |  | 944 | 70 | 1014 | 16.06 |
| Ephemeroptera | Ephemerellidae |  | 403 | 61 | 464 | 7.35 |
| Ephemeroptera | Ephemeridae |  | 20 | 318 | 338 | 5.35 |
| Ephemeroptera | Caenidae |  | 31 | 8 | 39 | 0.62 |
| Ephemeroptera | Ameletidae |  | 18 | 10 | 28 | 0.44 |
| Ephemeroptera | Baetidae |  | 6 | 1 | 7 | 0.11 |
| Plecoptera | Chloroperidae | *Alloperla ishikariana* | 102 | 130 | 232 | 3.67 |
| Plecoptera | Nemouridae |  | 111 | 16 | 127 | 2.01 |
| Plecoptera | Perlodidae | *Stavsolus ainu* | 41 | 3 | 44 | 0.70 |
| Plecoptera | Leuctridae |  | 0 | 15 | 15 | 0.24 |
| Plecoptera | Chloroperidae |  | 3 | 7 | 10 | 0.16 |
| Plecoptera | Capniidae |  | 6 | 0 | 6 | 0.10 |
| Trichoptera | Lepidostomatidae |  | 294 | 1045 | 1339 | 21.20 |
| Trichoptera | Leptoceridae |  | 192 | 193 | 385 | 6.10 |
| Trichoptera | Goeridae |  | 229 | 18 | 247 | 3.91 |
| Trichoptera | Hydropsychidae |  | 202 | 37 | 239 | 3.78 |
| Trichoptera | Stenopsychidae |  | 154 | 21 | 175 | 2.77 |
| Trichoptera | Apataniidae |  | 81 | 14 | 95 | 1.50 |
| Trichoptera | Brachycentridae |  | 90 | 2 | 92 | 1.46 |
| Trichoptera | Hydrobiosidae |  | 31 | 31 | 62 | 0.98 |
| Trichoptera | Polycentropodidae |  | 1 | 60 | 61 | 0.97 |
| Trichoptera | Glossosomatidae |  | 37 | 0 | 37 | 0.59 |
| Trichoptera | Rhyacophilidae |  | 30 | 6 | 36 | 0.57 |
| Trichoptera | Uenoiidae |  | 26 | 2 | 28 | 0.44 |
| Trichoptera | Limnephilidae |  | 2 | 5 | 7 | 0.11 |
| Trichoptera | Philopotamidae |  | 0 | 7 | 7 | 0.11 |
| Trichoptera | Molannidae |  | 0 | 6 | 6 | 0.10 |
| Trichoptera | Hydroptilidae |  | 1 | 0 | 1 | 0.02 |
