## Supplementary material for "Cryptic connectivity between hyporheic and riparian zones via winged aquatic insects revealed by DNA barcoding": SI7

**Supplementary information 7**

Sample information sequenced for mtDNA COI region in this study. HU: Hokkaido University; SU: Shinshu University. A list of sequences obtained from NCBI data base is also attached. All the own sequences are being in registration for DDBJ database and waiting for their accession numbers

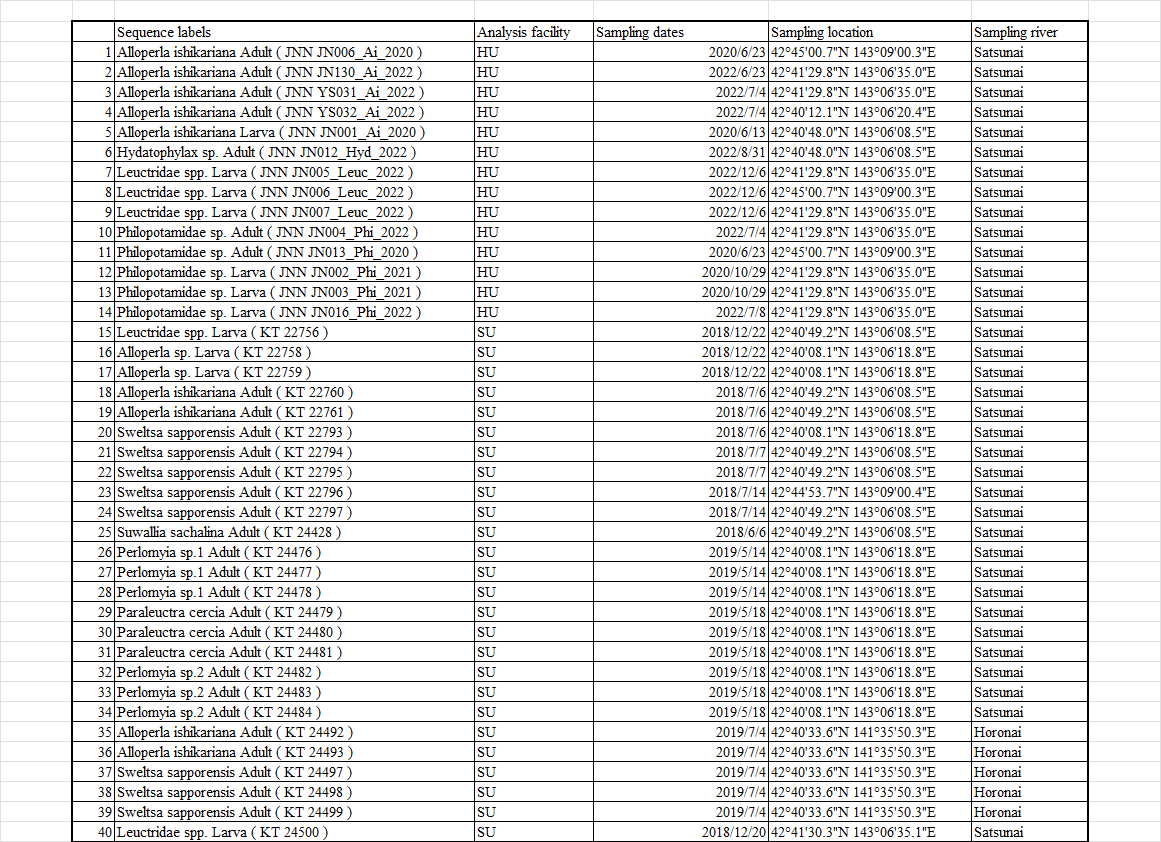

Continued

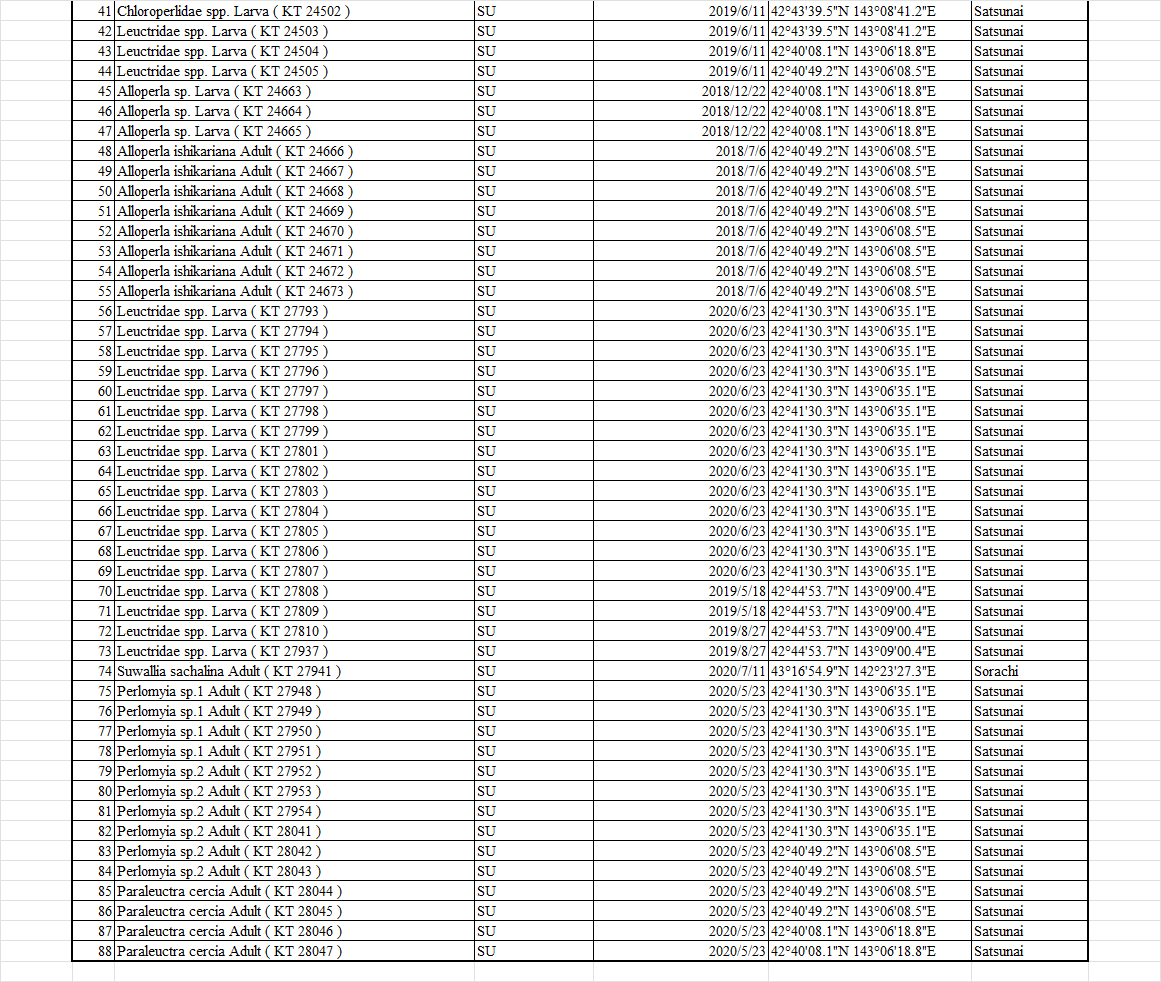

Continued

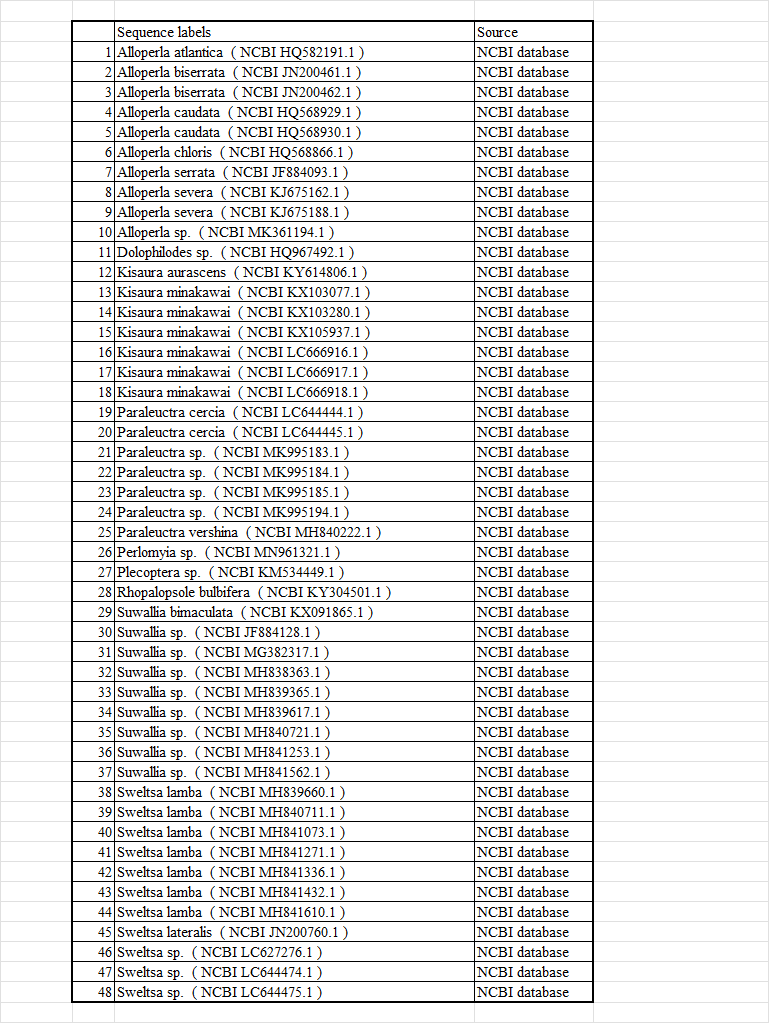
